# Muscle fiber type differences in nitrate and nitrite storage and nitric oxide signaling in rats

**DOI:** 10.1101/2020.06.01.128322

**Authors:** Gary M. Long, Derrick A. Gray, Ashley D. Troutman, Amanda Fisher, Mary Beth Brown, Andrew R. Coggan

## Abstract

Recent studies have emphasized the importance of the nitric oxide synthase (NOS)-independent, nitrate (NO_3_^−^) → nitrite (NO_2_^−^) → nitric oxide (NO) pathway in skeletal muscle. In particular, it has been hypothesized that this pathway is especially active in type II, or fast-twitch, muscle fibers, necessitating greater NO_3_^−^ and NO_2_^−^ storage. We therefore measured NO_3_^−^ and NO_2_^−^ concentrations in the predominantly fast-twitch vastus lateralis and predominantly slow-twitch soleus muscles of rats. Contrary to the above hypothesis, we found that NO_3_^−^ and NO_2_^−^ concentrations were 3.4-fold and 1.8-fold higher, respectively, in the soleus. On the other hand, NO signaling (i.e., cyclic guanosine monophosphate (cGMP) level) was comparable in the two muscles. Although the physiological significance of these observations remains to be determined, we speculate that NO production via the NO_3_^−^ → NO_2_^−^ → NO pathway is normally higher in slow-twitch muscles, thus helping compensate for their inherently lower NOS activity.

## Introduction

The gaseous free radical nitric oxide (NO) plays an important regulatory role in a wide variety of tissues. These include skeletal muscle, in which NO availability influences blood flow, glucose uptake, mitochondrial respiration, contractile function, etc. (1). These effects are mediated via both cyclic guanosine monophosphate (cGMP)-dependent and cGMP-independent mechanisms, e.g., S-nitrosylation of the ryanodine receptor (1). Originally, NO production in muscle (and other tissues) was believed to be entirely due to the activity of the enzyme NO synthase (NOS), which catalyzes the reaction: 2 L-arginine + 3 NADPH + 3 H^+^ + 4 O_2_ → 2 citrulline + 2 NO + 4 H_2_O + 3 NADP^+^. The NO thus produced can be eliminated via oxidation to first nitrite (NO_2_^−^) and then nitrate (NO_3_^−^), followed by urinary excretion. More recently, however, it has been recognized that NO can also be generated via reversal of this pathway, i.e., via the reduction of NO_3_^−^ to NO_2_^−^ and then to NO via bacterial and/or mammalian nitroreductases (e.g., xanthine oxidoreductase (XOR)) (2,3). Compared to NOS-mediated NO production, this alternative pathway tends to be favored at low pH and low pO_2_, conditions typical of resting and especially contracting muscle (4). In fact, due in part to its large mass skeletal muscle represents the body’s major NO_3_^−^ reservoir, which potentially can be used to support NO formation locally or, after export via the circulation, in other tissues (4). In turn, muscle seems to depend upon both endogenous and exogenous sources to maintain its stores of NO_3_^−^ and NO_2_^−^ (4,5).

Based in part on the effects of dietary NO_3_^−^ supplementation on muscle blood flow (6), it has been hypothesized that the NO_3_^−^ → NO_2_^−^ → NO pathway is especially active in type II, or fast-twitch, skeletal muscle (7). Accordingly, Nyakayiru, van Loon, and Verdijk (8) have recently speculated in this journal that NO_3_^−^ and NO_2_^−^ stores may be greater in fast-twitch muscle fibers. In contrast to this hypothesis, however, as reported herein we have found that NO_3_^−^ and NO_2_^−^ levels are actually several-fold higher in the predominantly type I, or slow-twitch, soleus vs. the predominantly fast-twitch superficial vastus lateralis of rats. To our knowledge, these are the first fiber type-specific measurements of muscle NO_3_^−^ and NO_2_^−^ concentrations in any species.

## Methods

Male Sprague Dawley rats (Charles River Laboratories, Wilmington, MA) weighing ~400g were heparinized, anesthetized (5% isofluorance in 95% O_2_), and 300 μL of blood withdrawn via the caudal artery. The sample was immediately centrifuged at 4°C for 10 min at 10,000 g to obtain plasma, which was stored at −80°C until analysis. Rats were then sacrificed via exsanguination and bilateral pneumectomy, after which sections of the soleus and the superficial (“white”) portion of the vastus lateralis were rapidly excised, snap-frozen in liquid N_2_, and also stored at −80°C until analysis. All study procedures were approved by the Institutional Animal Care and Use Committee at Indiana University (protocol number 11151).

Plasma and muscle NO_3_^−^ and NO_2_^−^ concentrations were measured using a high performance liquid chromatography (HPLC) analyzer (ENO-30, Eicom USA, San Diego, CA) as previously described in detail (9,10). Briefly, 25 μL of thawed plasma was mixed 1:1 with methanol, centrifuged at 4°C for 10 min at 10,000 g, and a 10 μL aliquot of the protein-poor supernatant injected into the HPLC. Muscle samples weighing ~10 mg were pulverized at liquid N_2_ temperature in a stainless steel tissue pulverizer (Bessman Tissue Pulverizer, Thermo Fischer Scientific), transferred into pre-weighed microcentrifuge tubes, and extracted on dry ice for 30 min using 50 μL methanol containing 0.5% Triton X-100 and 0.1 mmol/L oxypurinol (to block residual XOR activity). Samples were then quickly reweighed to determine the exact amount of tissue added, centrifuged at 4°C for 10 min at 10,000 g, and stored at − 80°C until analysis. A 10 μL aliquot of the tissue extract was subsequently injected into the HPLC. NO_3_^−^ and NO_2_^−^ levels in plasma and muscle were calculated from standard curves generated using NIST-traceable standard solutions.

Muscle cGMP content was measured using an ELISA kit (Cayman Chemicals, Ann Arbor, MI). A 25 μL aliquot of each tissue extract was transferred to the microplate, evaporated to dryness, and resuspended in 500 μL of assay buffer prior to analysis.

Normality of data distribution was tested using the D’Agostino-Pearson omnibus test. Differences between muscles were determined using unpaired *t* tests for unequal variance. P<0.05 was considered significant. Statistical analyses were performed using GraphPad Prism version 8.4.1 (GraphPad Software, La Jolla, CA).

## Results

As shown in Figure 1, muscle NO_3_^−^ and NO_2_^−^ concentrations were significantly higher (P<0.01) in the slow-twitch soleus than in the fast-twitch vastus lateralis. On the other hand, there was no difference in the cGMP content of the two muscles (Fig. 1).

**Figure 1.**
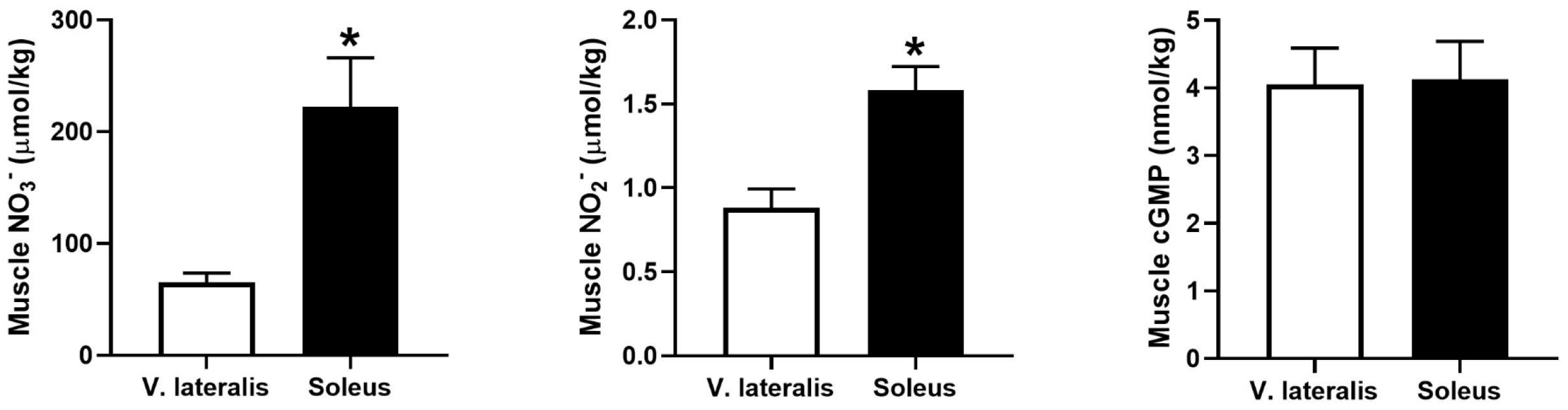
Nitrate (NO_3_^−^), nitrite (NO_2_^−^), and cyclic guanosine monophosphate (cGMP) concentrations in the superficial vastus lateralis and soleus muscles of male Sprague Dawley rats. *Soleus significantly higher than vastus lateralis (P<0.01). Values are mean ± S.E. for n = 9-10/group.

Muscle NO_3_^−^, NO_2_^−^, and/or cGMP concentrations were not significantly correlated with each other, in either muscle individually or when data from both muscles were combined (data not shown).

Plasma NO_3_^−^ and NO_2_^−^ concentrations averaged 29 ± 4 and 0.33 ± 0.04 μmol/L, respectively.

## Discussion

It has been known for over a decade that NO_3_^−^ and NO_2_^−^ levels are higher in various tissues (e.g., liver, heart) than in plasma (2,11–13). This includes both mouse (14) and human (15,16) skeletal muscle. However, interest in the latter tissue has recently increased (e.g., 17), due in part to studies by Piknova and co-workers (4,5,18,19) that have highlighted the central role of muscle in whole-body NO_3_^−^/NO_2_^−^/NO metabolism. Skeletal muscle, though, is a heterogeneous tissue, composed of two major fiber types that differ significantly not only in terms of their contractile properties, blood supply, energy metabolism, etc., but also in terms of NOS-mediated NO signaling (see below). In light of recent publications emphasizing the importance of NO production via the alternative NO_3_^−^ → NO_2_^−^ → NO pathway in muscle, we therefore sought to determine whether there are also fiber type differences in NO_3_^−^ and NO_2_^−^ storage. To do so, we used a highly-sensitive HPLC-based method (10) to measure NO_3_^−^ and NO_2_^−^ concentrations in the superficial vastus lateralis and soleus muscles of rats. These hindlimb muscles were chosen due to their extremes of fiber type distribution, being composed of approximately 97% fast-twitch and 87% slow-twitch fibers, respectively (20). We found that NO_3_^−^ and NO_2_^−^ concentrations were 3.4- and 1.8-fold higher, respectively, in the predominantly slow-twitch soleus than in the predominantly fast-twitch vastus lateralis. Extrapolating to muscles theoretically composed of 100% of just one fiber type, this implies differences in NO_3_^−^ and NO_2_^−^ content between slow- and fast-twitch fibers of ~4-fold and ~2-fold, respectively. Our data therefore do not support the hypothesis of Nyakayiru, van Loon, and Verdijk (8) that NO_3_^−^ and NO_2_^−^ storage are greater in fast-twitch fibers, at least in rats.

As indicated previously, it is well-established that, at least in rodents, NOS-dependent NO signaling is more prominent in fast-twitch than in slow-twitch muscle. For example, expression of NOS1, or nNOS, is greatest in rat fast-twitch fibers, and minimal or absent in slow-twitch muscle fibers (1). Correspondingly, total NOS activity is higher in muscles composed primarily of fast-twitch fibers (e.g., extensor digitorum longus, gastrocnemius, plantaris) than in the primarily slow-twitch soleus (1). NOS activators or inhibitors (e.g., L-NAME) also generally exhibit far greater effects on, e.g., contractile function in fast-twitch vs. slow-twitch muscle (1). Despite this, it does not necessarily follow that NOS-independent NO signaling is also greater in fast-twitch muscle. Indeed, one could argue that limited or absent NOS activity in rodent slow-twitch fibers would require more dependence upon the NO_3_^−^ → NO_2_^−^ → NO pathway to augment or enable NO production. If so, this would likely require greater storage of NO_3_^−^/NO_2_^−^ in slow-twitch muscle, especially since such fibers are more readily recruited, and thus more NO_3_^−^ and NO_2_^−^ presumably regularly utilized, during ordinary contractile activity (e.g., stance, ambulation). Furthermore, despite lower microvascular pO_2_ in fast-twitch vs. slow-twitch muscle at rest (i.e., ~25 vs. ~30 mmHg) and especially during electrically-stimulated contractions (i.e., ~10 vs. ~25 mmHg), which is hypothesized to enhance deoxyhemoglobin-mediated reduction of NO_2_^−^ to NO in the capillaries (6), the rate of intracellular reduction of NO_2_^−^ to NO may actually normally be greater, or at least equal, in slow-twitch muscle. This is because 1) the bimolecular rate constant for NO_2_^−^ reduction by myoglobin is twice that of hemoglobin, such that in, e.g., the heart most NO_2_^−^ reduction actually occurs in the myocytes, not in the circulation (11), and 2) the total concentration of myoglobin is several-fold higher in slow-twitch muscle (21). Due to the hyperbolic shape and low p50 of the oxymyoglobin dissociation curve, the latter difference means that the deoxyhemoglobin concentration will also generally be higher in slow-twitch muscle – only at very low intracellular pO_2_, such as may occur during ischemia or intense exercise (22,23), would this not be true. Reduction of NO_2_^−^ to NO (or NO_3_^−^ to NO_2_^−^) by XOR may also very well be higher in slow-twitch muscle. This is because numerous immunohistochemical studies have demonstrated that XOR expression in muscle tissue is limited to vascular endothelial cells (24–27), and capillarization is much greater in slow-twitch muscle. Consistent with the possibility of significant NO production via the NO_3_^−^ → NO_2_^−^ → NO pathway in slow-twitch fibers, we found that, at least in the basal state, the concentration of the canonical NO second messenger cGMP is just as high in the soleus as the vastus lateralis, despite limited or absent NOS activity in rodent slow-twitch muscle.

Along with the available literature (e.g., 1,3,11,21,28–33), we therefore interpret our novel observation that NO_3_^−^ and NO_2_^−^ concentrations are higher in the soleus than the superficial vastus lateralis muscle of rats as suggesting that although NOS-dependent NO production and signaling are more prominent in fast-twitch fibers, the NOS-independent, NO_3_^−^ → NO_2_^−^ → NO pathway may be favored in slow-twitch fibers. This hypothesis is shown in Figure 2, which also illustrates potential differences in the mechanisms by which NO exerts its physiological effects in the two fiber types (i.e., by cGMP-dependent vs. cGMP-independent pathways). Although speculative, we believe that at a minimum this schema can serve as a useful “roadmap” for future research. For example, it would be of interest to quantify XOR activity in both slow-twitch and fast-twitch muscles, which to our knowledge has not been done. Similarly, the higher NO_3_^−^ and NO_2_^−^ concentrations we found in the soleus implies a higher rate of import of these ions by slow-twitch fibers. It would therefore be of interest to determine whether expression of the NO_3_^−^/H^+^ antiporter sialin, present in both pig (34) and human (35) skeletal muscle, is higher in slow-twitch vs. fast-twitch fibers.

**Figure 2.**
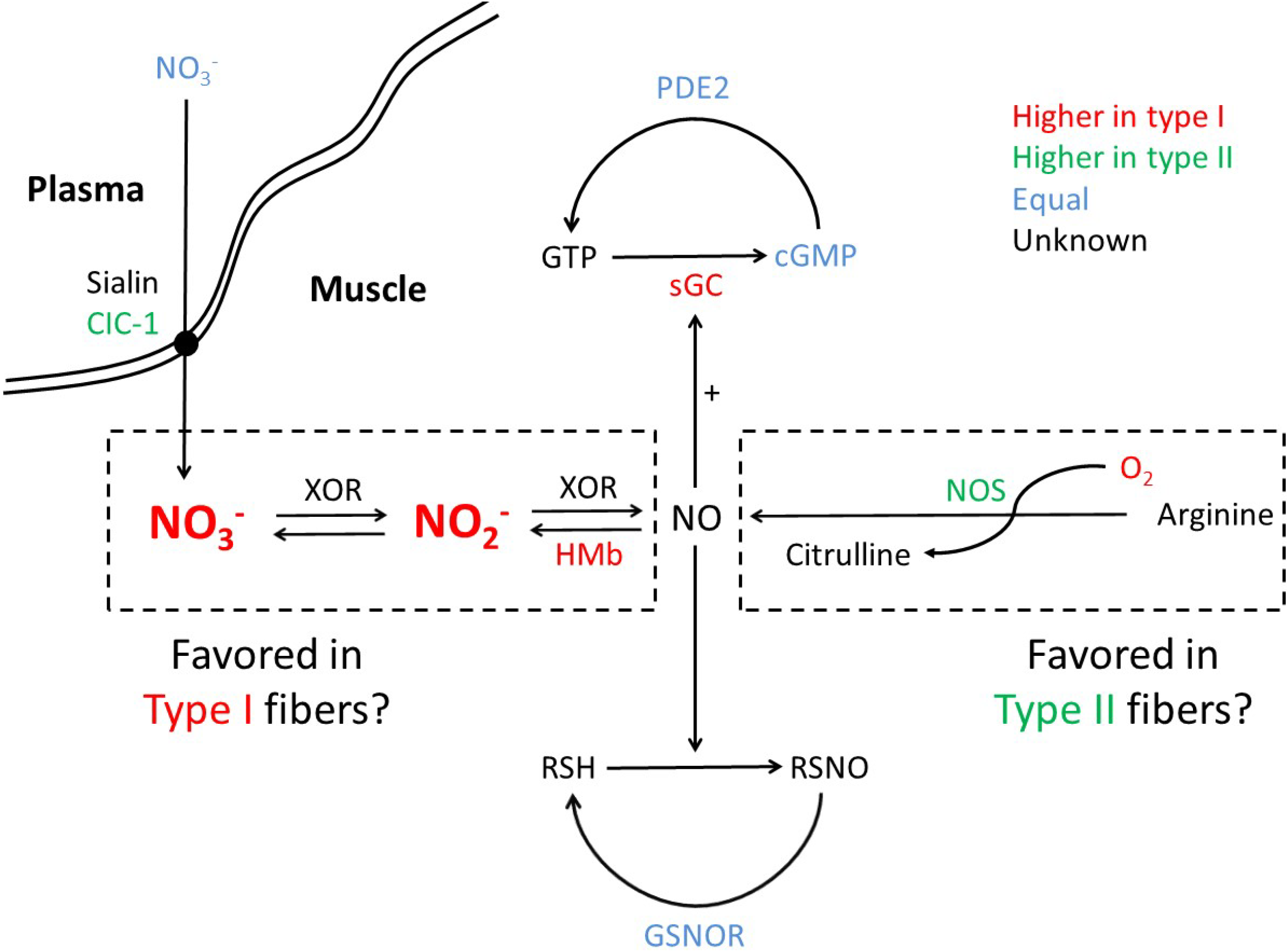
Proposed schema of NO signaling in slow-twitch (type I) and fast-twitch (type II) muscle fibers, based on the present results along with the published literature (Refs. 1,3,11,21,28–33). Abbreviations: ClC-1, chloride channel 1. cGMP, cyclic guanosine monophosphate. GTP, guanosine triphosphate. GSNOR, *S*-nitrosoglutahione reductase. HMb, deoxymyoglobin. NOS, nitric oxide synthase. PDE2, phosphodiesterase 2. RSH, thiol groups. RSNO, S-nitrosothiols. sGC, soluble guanyl cyclase. XOR, xanthine oxidoreductase. See Discussion for additional details.

As with any study, there are limitations to the present research. We only measured NO_3_^−^, NO_2_^−^, and cGMP levels, and did not quantify any of the other metabolites or proteins shown in Figure 2. We also made measurements only in resting muscle, and did not evaluate the effects of, e.g., increased contractile activity. Thus, although supported by the available literature our hypothesis that NO production via the NO_3_^−^ → NO_2_^−^ → NO pathway is normally greater in slow-twitch than in fast-twitch muscle is clearly still untested. We also did not investigate the effects of dietary NO_2_^−^ supplementation, which despite the differences in muscle NO_3_^−^ and NO_2_^−^ that we observed following a normal diet could still preferentially “target” fast-twitch muscle (7). Finally, we studied rats, and the present results need to be replicated in humans, especially given marked differences between rats and humans in other aspects of NO_3_^−^/NO_2_^−^/NO metabolism, especially fiber type differences in NOS expression (1). The present method is theoretically sensitive enough to quantify the NO_3_^−^ concentration of individual freeze-dried human muscle fibers from biopsy samples, but it would require pooling ~100 fibers of a given type to also measure NO_2_^−^ concentration.

In summary, we have found that, contrary to previous suggestions, NO_3_^−^ and NO_2_^−^ concentrations are markedly higher in the predominantly slow-twitch soleus vs. the predominantly fast-twitch superficial vastus lateralis of rats. These are the first fiber type-specific measurements of these NO precursors in any species. The physiological significance of this difference remains to be determined.

## Acknowledgements

This publication was made possible by award numbers HL121661 from the National Heart, Lung, and Blood Institute (NHLBI) and AG053606 from the National Institute on Aging (NIA) of the National Institutes of Health (NIH). Its contents are solely the responsibility of the authors and do not necessarily represent the official views of the NHLBI, NIA, or NIH.

## Notes

### Competing Interest Statement

The authors have declared no competing interest.

